## Supplementary Figure S1 for "Development of a versatile high-throughput mutagenesis assay with multiplexed short read NGS using DNA-barcoded *supF* shuttle vector library amplified in non-SOS *E. coli*"

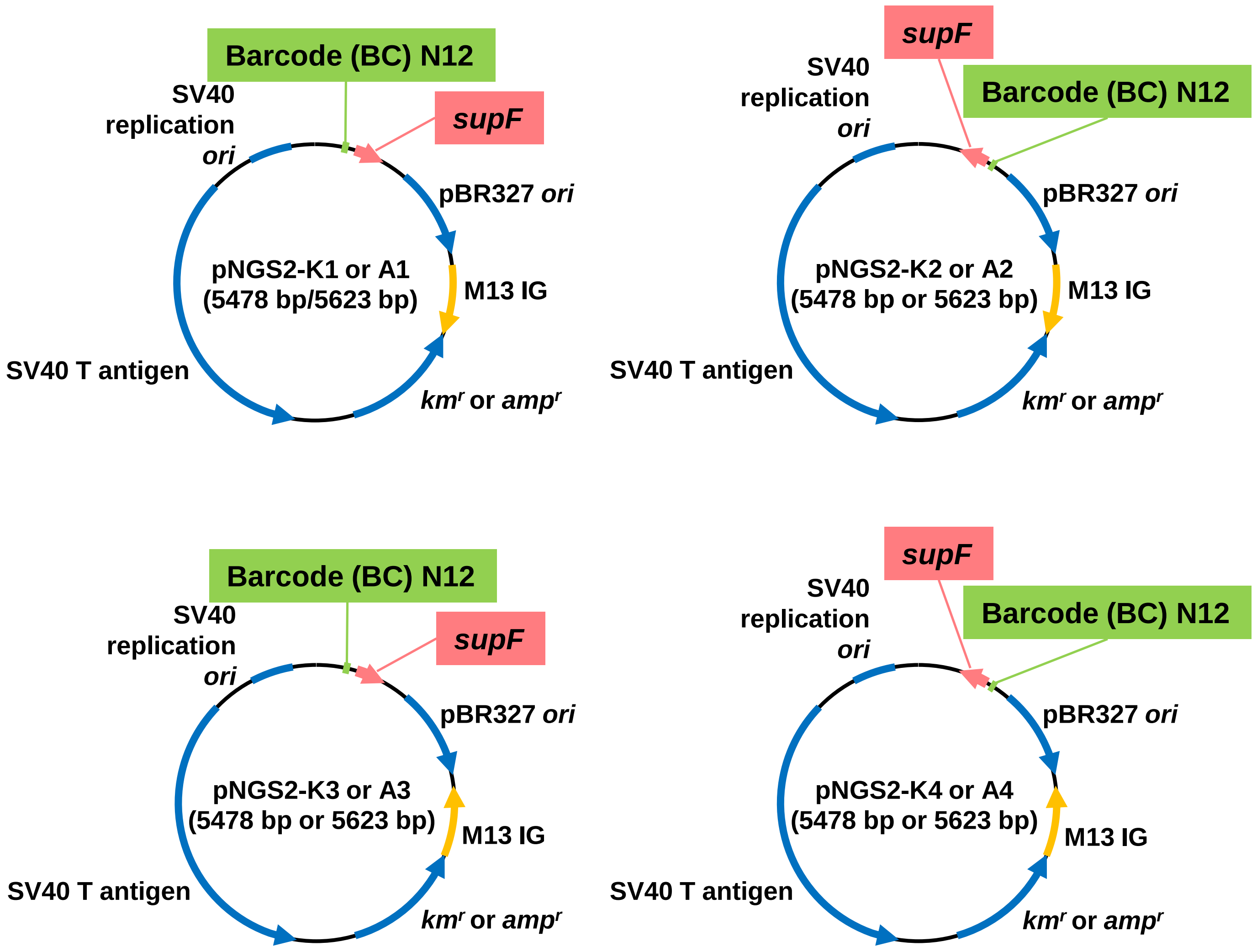


**Supplementary Figure S1. *SupF* shuttle vector $N_{12}$-BC library**
The circular maps of eight series of *supF* shuttle vector plasmids, pNGS2 -K1, -K2, -K3, -K4, -A1, -A2, -A3, -A4. Each plasmid vector contains an amber suppressor tRNA gene (*supF*), a TP53-/Rb- binding-deficient mutant SV40 large T antigen (E107K/D402E) with a SV40 replication origin, a pBR327 origin of replication, a M13 intergenic region, and either an ampicillin-resistance gene (*amp*$^r$) or kanamycin-resistance gene (*km*$^r$) in the indicated direction.
