## Supplementary Figure S2 for "Development of a versatile high-throughput mutagenesis assay with multiplexed short read NGS using DNA-barcoded *supF* shuttle vector library amplified in non-SOS *E. coli*"

| Primer Name (Forward $N_0$) | Sequence (5' to 3') |
|---|---|
| BC12-ID01-F | ATGCAATTAATTGTAAAGGCCTCAGCGAATT |
| BC12-ID02-F | ATGCAAGGAATTGTAAAGGCCTCAGCGAATT |
| BC12-ID03-F | ATGCAACCAATTGTAAAGGCCTCAGCGAATT |
| BC12-ID04-F | ATGCTTAATTTTGTAAAGGCCTCAGCGAATT |
| BC12-ID05-F | ATGCTTGGTTTTGTAAAGGCCTCAGCGAATT |
| BC12-ID06-F | ATGCTTCCTTTTGTAAAGGCCTCAGCGAATT |
| BC12-ID07-F | ATGCGGAAGGTTGTAAAGGCCTCAGCGAATT |
| BC12-ID08-F | ATGCGGTTGGTTGTAAAGGCCTCAGCGAATT |
| BC12-ID09-F | ATGCGGCCGGTTGTAAAGGCCTCAGCGAATT |
| BC12-ID10-F | ATGCCCAACCTTGTAAAGGCCTCAGCGAATT |
| BC12-ID11-F | ATGCCCTTCCTTGTAAAGGCCTCAGCGAATT |
| BC12-ID12-F | ATGCCCGGCCTTGTAAAGGCCTCAGCGAATT |
| BC12-ID13-F | ATGCAATTGGTTGTAAAGGCCTCAGCGAATT |
| BC12-ID14-F | ATGCAAGGTTTTGTAAAGGCCTCAGCGAATT |
| BC12-ID15-F | ATGCTTAAGGTTGTAAAGGCCTCAGCGAATT |
| BC12-ID16-F | ATGCTTGGAATTGTAAAGGCCTCAGCGAATT |
| BC12-ID17-F | ATGCGGAATTTTGTAAAGGCCTCAGCGAATT |
| BC12-ID18-F | ATGCGGTTAATTGTAAAGGCCTCAGCGAATT |
| BC12-ID19-F | ATGCAATTCCTTGTAAAGGCCTCAGCGAATT |
| BC12-ID20-F | ATGCAACCTTTTGTAAAGGCCTCAGCGAATT |
| BC12-ID21-F | ATGCTTAACCTTGTAAAGGCCTCAGCGAATT |
| BC12-ID22-F | ATGCTTCCAATTGTAAAGGCCTCAGCGAATT |
| BC12-ID23-F | ATGCCCAATTTTGTAAAGGCCTCAGCGAATT |
| BC12-ID24-F | ATGCCCTTAATTGTAAAGGCCTCAGCGAATT |

| Primer Name (Forward $N_2$) | Sequence (5' to 3') |
|---|---|
| BC12-ID49-NN-F | NNATGCATGATGTTGTAAAGGCCTCAGCGAATT |
| BC12-ID50-NN-F | NNATGCAGTAGTTTGTAAAGGCCTCAGCGAATT |
| BC12-ID51-NN-F | NNATGCTAGTAGTTGTAAAGGCCTCAGCGAATT |
| BC12-ID52-NN-F | NNATGCTGATGATTGTAAAGGCCTCAGCGAATT |
| BC12-ID53-NN-F | NNATGCGATGATTTGTAAAGGCCTCAGCGAATT |
| BC12-ID54-NN-F | NNATGCGTAGTATTGTAAAGGCCTCAGCGAATT |
| BC12-ID55-NN-F | NNATGCATCATCTTGTAAAGGCCTCAGCGAATT |
| BC12-ID56-NN-F | NNATGCACTACTTTGTAAAGGCCTCAGCGAATT |
| BC12-ID57-NN-F | NNATGCTACTACTTGTAAAGGCCTCAGCGAATT |
| BC12-ID58-NN-F | NNATGCTCATCATTGTAAAGGCCTCAGCGAATT |
| BC12-ID59-NN-F | NNATGCCATCATTTGTAAAGGCCTCAGCGAATT |
| BC12-ID60-NN-F | NNATGCCTACTATTGTAAAGGCCTCAGCGAATT |
| BC12-ID61-NN-F | NNATGCAGCAGCTTGTAAAGGCCTCAGCGAATT |
| BC12-ID62-NN-F | NNATGCACGACGTTGTAAAGGCCTCAGCGAATT |
| BC12-ID63-NN-F | NNATGCGACGACTTGTAAAGGCCTCAGCGAATT |
| BC12-ID64-NN-F | NNATGCGCAGCATTGTAAAGGCCTCAGCGAATT |
| BC12-ID65-NN-F | NNATGCCAGCAGTTGTAAAGGCCTCAGCGAATT |
| BC12-ID66-NN-F | NNATGCCGACGATTGTAAAGGCCTCAGCGAATT |
| BC12-ID67-NN-F | NNATGCTGCTGCTTGTAAAGGCCTCAGCGAATT |
| BC12-ID68-NN-F | NNATGCTCGTCGTTGTAAAGGCCTCAGCGAATT |
| BC12-ID69-NN-F | NNATGCGTCGTCTTGTAAAGGCCTCAGCGAATT |
| BC12-ID70-NN-F | NNATGCGCTGCTTTGTAAAGGCCTCAGCGAATT |
| BC12-ID71-NN-F | NNATGCCTGCTGTTGTAAAGGCCTCAGCGAATT |
| BC12-ID72-NN-F | NNATGCCGTCGTTTGTAAAGGCCTCAGCGAATT |

| Primer Name (Forward $N_1$) | Sequence (5' to 3') |
|---|---|
| BC12-ID25-N-F | NATGCAAGGCCTTGTAAAGGCCTCAGCGAATT |
| BC12-ID26-N-F | NATGCAACCGGTTGTAAAGGCCTCAGCGAATT |
| BC12-ID27-N-F | NATGCGGAACCTTGTAAAGGCCTCAGCGAATT |
| BC12-ID28-N-F | NATGCGGCCAATTGTAAAGGCCTCAGCGAATT |
| BC12-ID29-N-F | NATGCCCAAGGTTGTAAAGGCCTCAGCGAATT |
| BC12-ID30-N-F | NATGCCCGGAATTGTAAAGGCCTCAGCGAATT |
| BC12-ID31-N-F | NATGCTTGGCCTTGTAAAGGCCTCAGCGAATT |
| BC12-ID32-N-F | NATGCTTCCGGTTGTAAAGGCCTCAGCGAATT |
| BC12-ID33-N-F | NATGCGGTTCCTTGTAAAGGCCTCAGCGAATT |
| BC12-ID34-N-F | NATGCGGCCTTTTGTAAAGGCCTCAGCGAATT |
| BC12-ID35-N-F | NATGCCCTTGGTTGTAAAGGCCTCAGCGAATT |
| BC12-ID36-N-F | NATGCCCGGTTTTGTAAAGGCCTCAGCGAATT |
| BC12-ID37-N-F | NATGCATAATATTGTAAAGGCCTCAGCGAATT |
| BC12-ID38-N-F | NATGCAGAAGATTGTAAAGGCCTCAGCGAATT |
| BC12-ID39-N-F | NATGCACAACATTGTAAAGGCCTCAGCGAATT |
| BC12-ID40-N-F | NATGCTATTATTTGTAAAGGCCTCAGCGAATT |
| BC12-ID41-N-F | NATGCTGTTGTTTGTAAAGGCCTCAGCGAATT |
| BC12-ID42-N-F | NATGCTCTTCTTTGTAAAGGCCTCAGCGAATT |
| BC12-ID43-N-F | NATGCGAGGAGTTGTAAAGGCCTCAGCGAATT |
| BC12-ID44-N-F | NATGCGTGGTGTTGTAAAGGCCTCAGCGAATT |
| BC12-ID45-N-F | NATGCGCGGCGTTGTAAAGGCCTCAGCGAATT |
| BC12-ID46-N-F | NATGCCACCACTTGTAAAGGCCTCAGCGAATT |
| BC12-ID47-N-F | NATGCCTCCTCTTGTAAAGGCCTCAGCGAATT |
| BC12-ID48-N-F | NATGCCGCCGCTTGTAAAGGCCTCAGCGAATT |

| Primer Name (Reverse $N_{0\sim2}$) | Sequence (5' to 3') |
|---|---|
| BC12-$N_0$-R | CTAACCAAGTTCCTCTTTCAG |
| BC12-$N_1$-R | NCTAACCAAGTTCCTCTTTCAG |
| BC12-$N_2$-R | NNCTAAACCAAGTTCCTCTTTCAG |

**Supplementary Figure S2. The nucleotide sequences of primer sets for multiplexing sample preparation for NGS**

The each $N_{12}$-BC library was independently amplified by PCR with a forward primer (BC12-IDXX-$N_X$-F) and a reverse primer (BC12-$N_X$-R). Pre-designed 6-nucleotide sequence (N6) were shown in red color.
