## Supplementary Figure S3 for "Development of a versatile high-throughput mutagenesis assay with multiplexed short read NGS using DNA-barcoded *supF* shuttle vector library amplified in non-SOS *E. coli*"

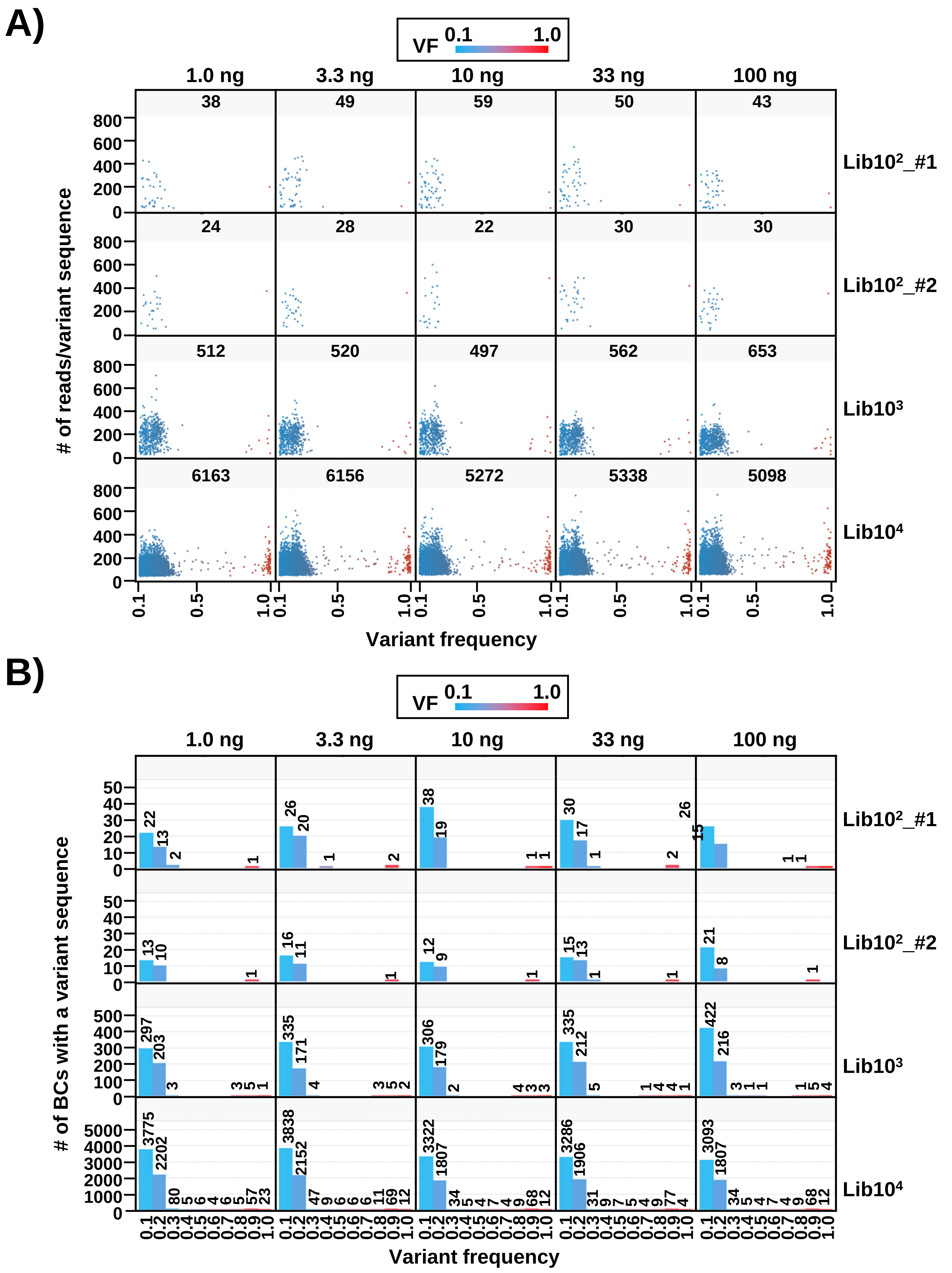


**Supplementary Figure S3. Variant frequency in $N_{12}$-BC sequences from the multiplexed NGS libraries**
(A) Dot plots of the distribution of the number of reads for each variant sequence according to their variant frequency for the samples in Fig. 2. The number in each segment represents the total number of $N_{12}$-BC sequences with a variant sequence. Source data is available in Supplementary Table S2. (B) Distribution of the number of $N_{12}$-BC sequences with a variant by variant frequency, shown as histograms with bin width 0.1 of the VF.
