## Supplementary Figure S7 for "Development of a versatile high-throughput mutagenesis assay with multiplexed short read NGS using DNA-barcoded *supF* shuttle vector library amplified in non-SOS *E. coli*"

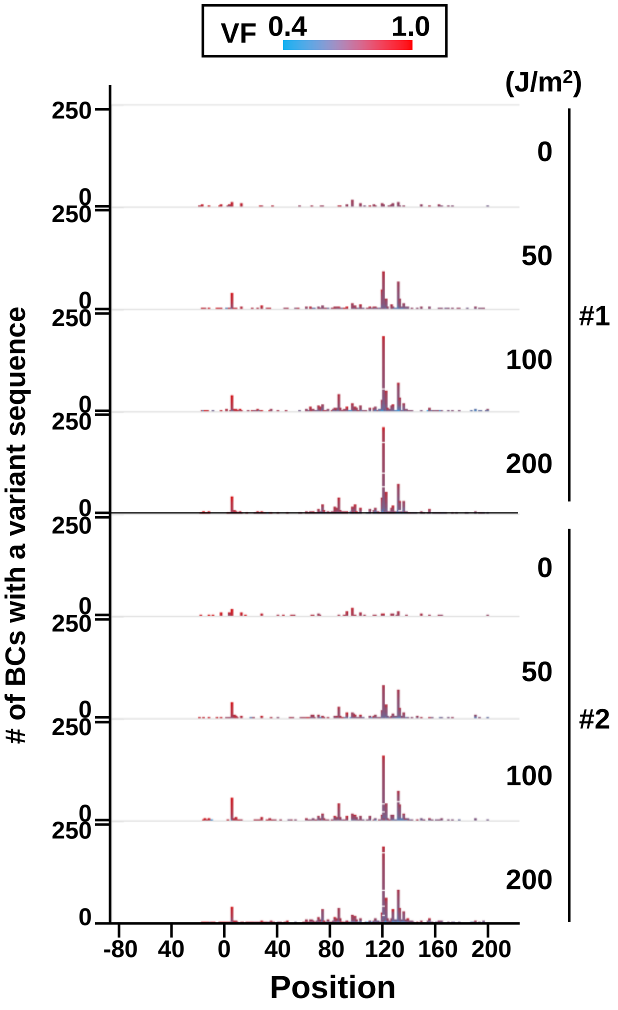


**Supplementary Figure S7. Number of $N_{12}$-BC sequences with a variant exceeding VF 0.4 at different UV-C doses and their distribution by nucleotide position (in mammalian cells, data from selection plates).** The colors reflect the variant frequency as indicated by the heatmap on top.
