## Supplementary Figure S8 for "Development of a versatile high-throughput mutagenesis assay with multiplexed short read NGS using DNA-barcoded *supF* shuttle vector library amplified in non-SOS *E. coli*"

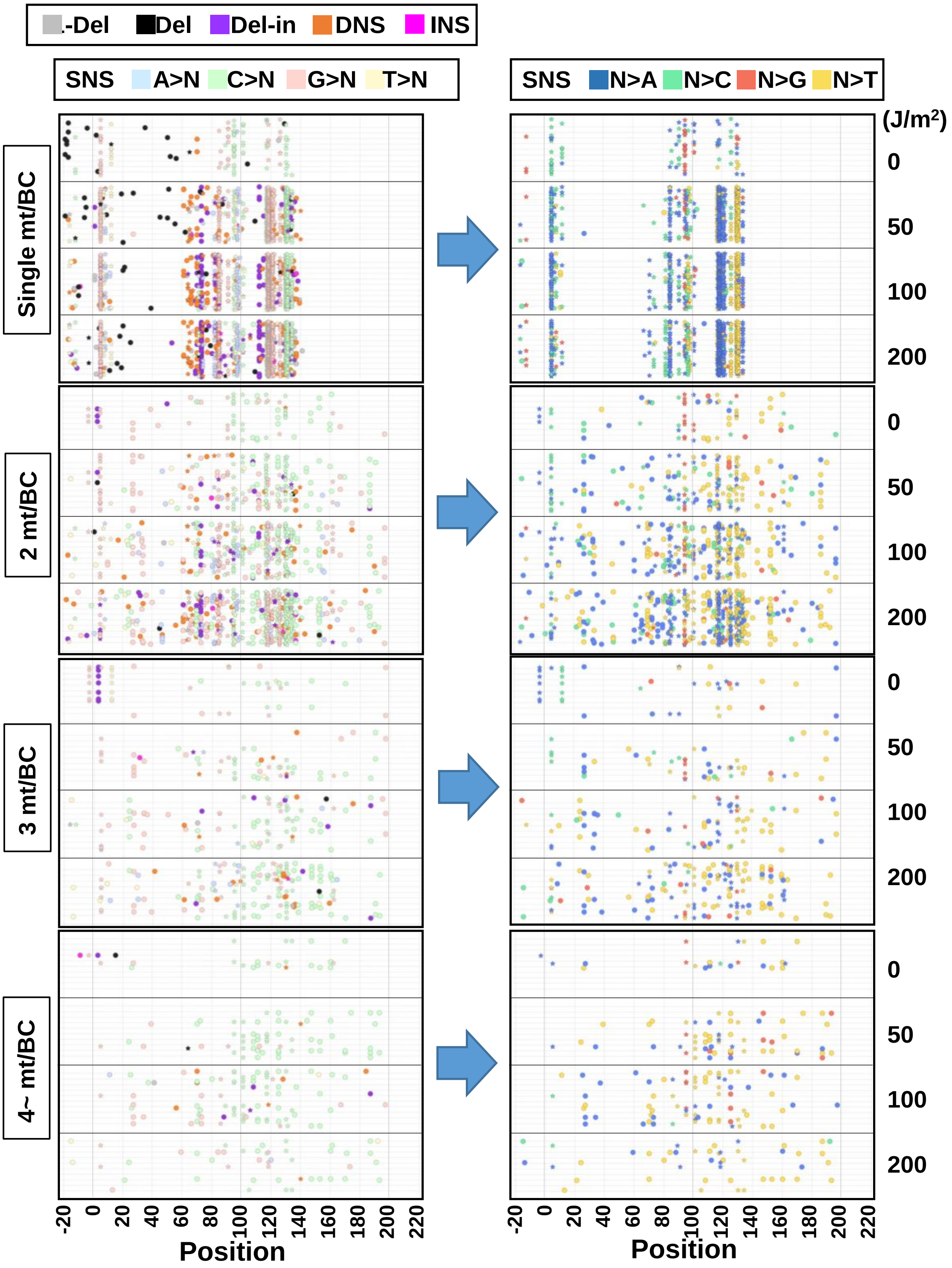


**Supplementary Figure S8. The position and mutation type of each identified mutation (data from selection plates, 0-200 J/m$^2$)**

The position and mutation type of each identified mutation are presented in dot plots for the cases of 1, 2, 3, or 4~ mutations per BC. The star symbols indicate the positions of mutations associated with mutant phenotype of the *supF* gene. Each type of mutation is indicated by a different color in the left-side plots (legend on top). The bases after the substitution are indicated by a different color for only SNSs in the right-side plots (legend on top). The mutations in identical BCs are plotted on the same horizontal line. Data is combined from Lib10$^4$_#1 and Lib10$^4$_#2.
