## Supplementary Figure S9 for "Development of a versatile high-throughput mutagenesis assay with multiplexed short read NGS using DNA-barcoded *supF* shuttle vector library amplified in non-SOS *E. coli*"

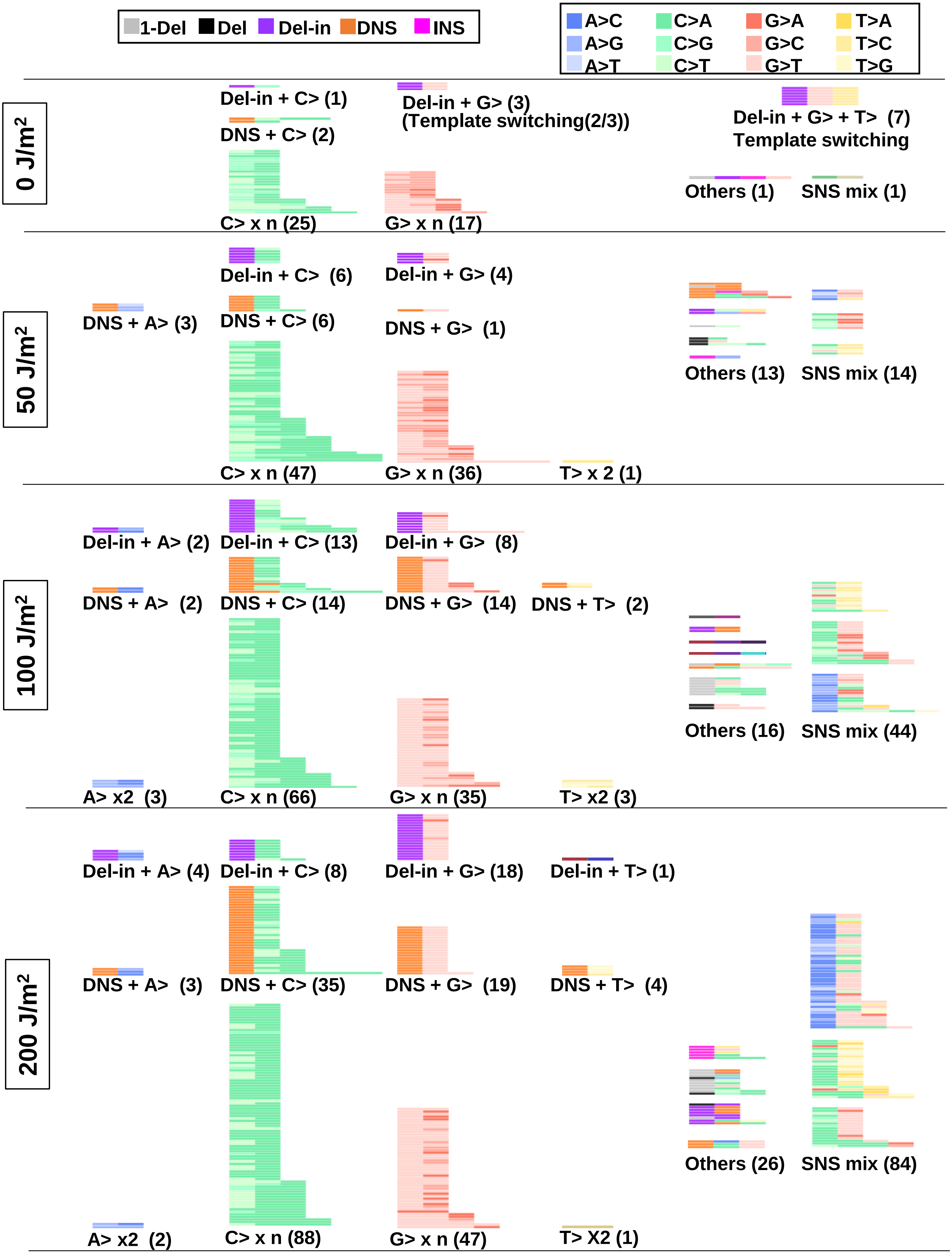


**Supplementary Figure S9. The combinations of UV-induced mutation types in the *supF* gene in mammalian cells (data from selection plates, 0-200 J/m$^2$)**

Combinations of mutation sequences for each BC with multiple mutations for each UV-dose. Combinations of more than one mutation per BC are shown. Each type of mutation is indicated by a different color and the type of aberration. The number in brackets and the height of stacked horizontal bars represent the number of BCs with the mutation(s).
