## Supplementary Figure S10 for "Development of a versatile high-throughput mutagenesis assay with multiplexed short read NGS using DNA-barcoded *supF* shuttle vector library amplified in non-SOS *E. coli*"

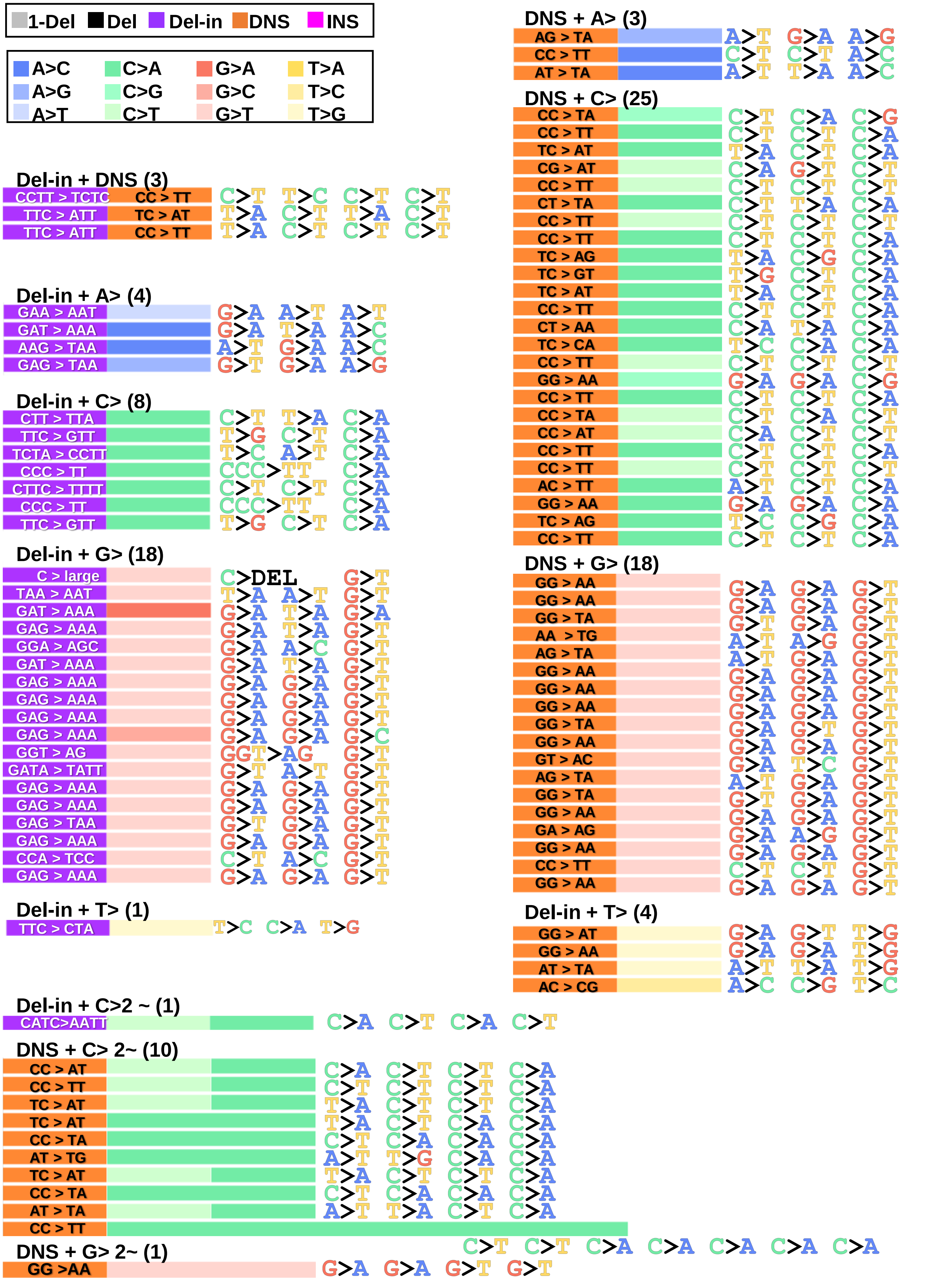


**Supplementary Figure S10. The combinations of UV-induced mutation types and mutation spectrum in the *supF* gene in mammalian cells (data from selection plates, 200 J/m²)**

Combinations of mutation sequences for each BC with multiple mutations for 200 J/m². Each type of mutation is indicated by a different color (legend on top). The horizontal bars reflect the type, sequence and combinations of mutations, and the number of combinations is indicated next to each categorized group of horizontal bars.
