## Supplementary Figure S11 for "Development of a versatile high-throughput mutagenesis assay with multiplexed short read NGS using DNA-barcoded *supF* shuttle vector library amplified in non-SOS *E. coli*"

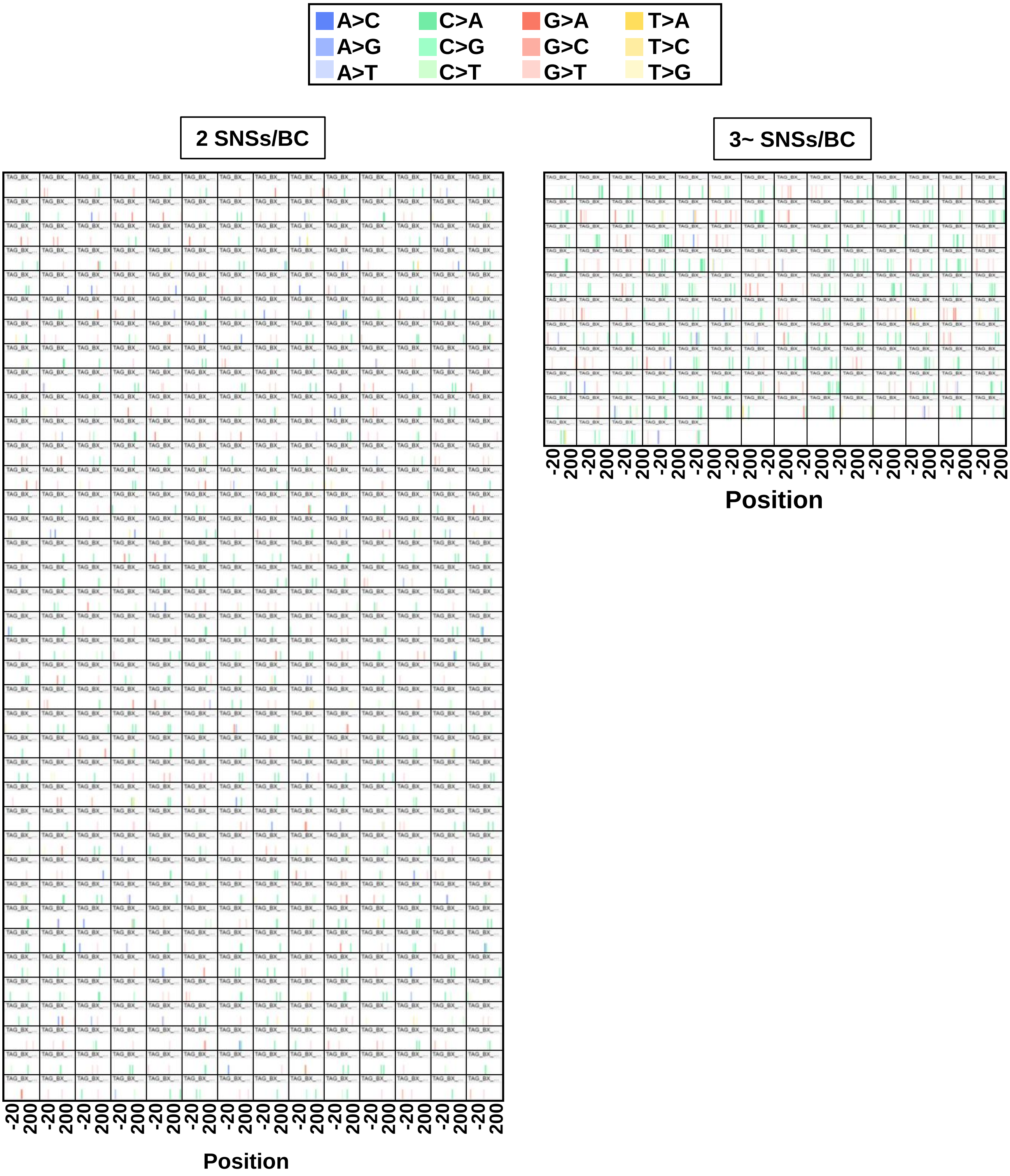


**Supplementary Figure S11. The distance between multiple SNSs per BC (data from selection plates, 50, 100, and 200 J/m$^2$)**

The position and type of each identified mutation per BC are presented in bar graphs for the cases of 2 (left-side graph) or 3~ (right-side graph) mutations per BC (233 bp, positions -19 to 214). Each single base substitution is indicated by a different color (legend on top). Data is combined from Lib10$^4$_#1 and Lib10$^4$_#2.
