## Supplementary Figure S12 for "Development of a versatile high-throughput mutagenesis assay with multiplexed short read NGS using DNA-barcoded *supF* shuttle vector library amplified in non-SOS *E. coli*"

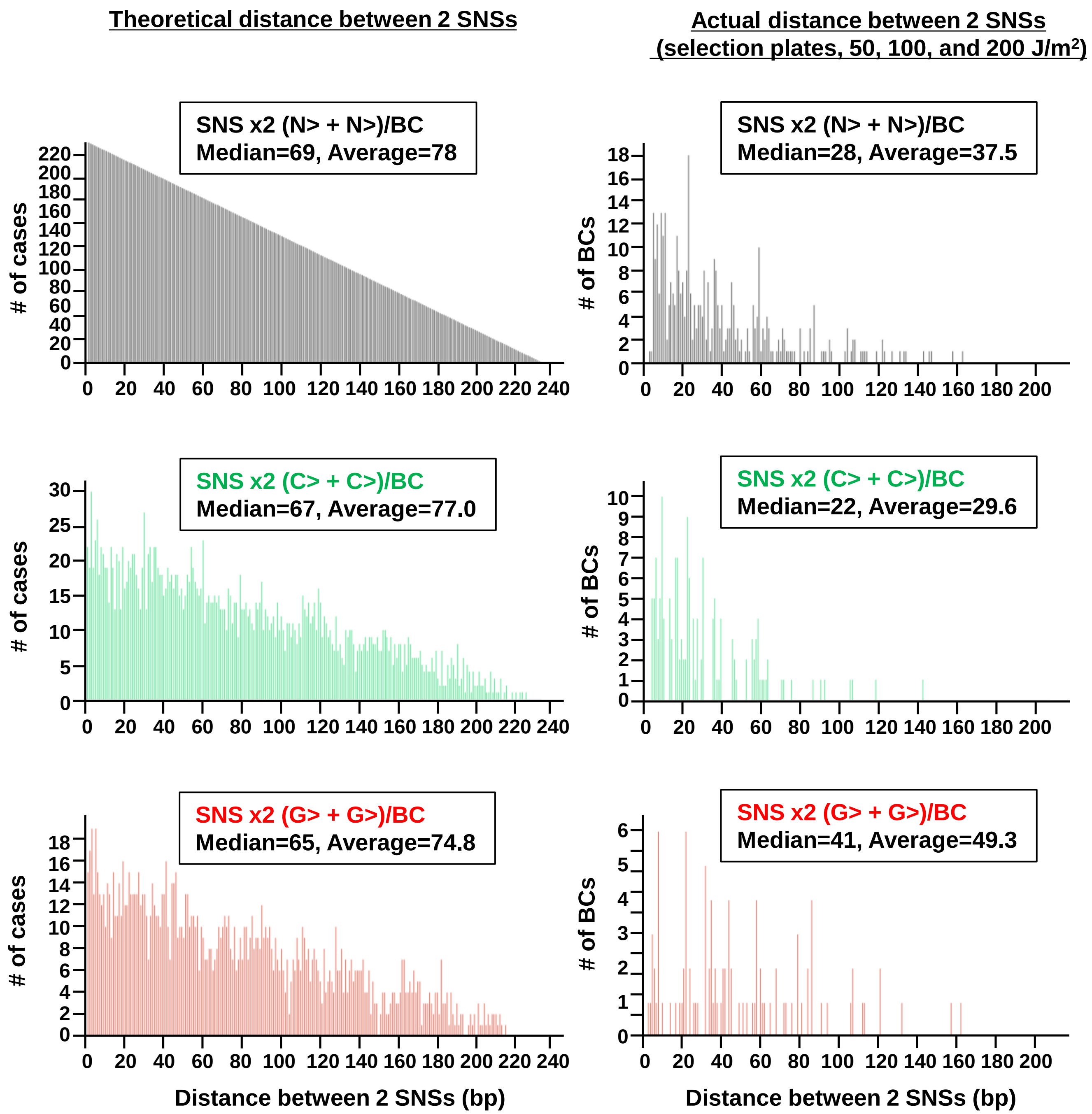


**Supplementary Figure S12. The distribution of the distance between 2 SNSs per BC (data from selection plates, 50, 100, and 200 J/m$^2$)**

The theoretical distances (bp) between 2 bases based on the sequence (233 bp, positions -19 to 214) are shown in the left-side histograms for all four bases (top), for two cytosines (middle), and for two guanines (bottom). The actual distances between 2 SNSs induced by UV per BC are shown on the right side. Data is combined from Lib10$^4$_#1 and Lib10$^4$_#2. The median and average distance are shown on the top of each histogram.
